# Human walking in the real world: Interactions between terrain type, gait parameters, and energy expenditure

**DOI:** 10.1101/2019.12.29.890434

**Authors:** DB Kowalsky, JR Rebula, LV Ojeda, PG Adamczyk, AD Kuo

**Author notes:** A. D. Kuo, corresponding author present address, Faculty of Kinesiology & Program in Biomedical Engineering, University of Calgary, Calgary, Alberta, CANADA.

## Abstract

Humans often traverse real-world environments with a variety of surface irregularities and inconsistencies, which can disrupt steady gait and require additional effort. Such effects have, however, scarcely been demonstrated quantitatively, because few laboratory biomechanical measures apply outdoors. Walking can nevertheless be quantified by other means. In particular, the foot’s trajectory in space can be reconstructed from foot-mounted inertial measurement units (IMUs), to yield measures of stride and associated variabilities. But it remains unknown whether such measures are related to metabolic energy expenditure. We therefore quantified the effect of five different outdoor terrains on foot motion (from IMUs) and net metabolic rate (from oxygen consumption) in healthy adults (N = 10; walking at 1.25 m/s). Energy expenditure increased significantly (P < 0.05) in the order Sidewalk, Dirt, Gravel, Grass, and Woodchips, with Woodchips about 27% costlier than Sidewalk. Terrain type also affected measures, particularly stride variability and virtual foot clearance (swing foot’s lowest height above consecutive footfalls). In combination, such measures can also roughly predict metabolic cost (adjusted *R*^2^ = 0.52, partial least squares regression), and even discriminate between terrain types (10% reclassification error). Body-worn sensors can characterize how uneven terrain affects gait, gait variability, and metabolic cost in the real world.

## Introduction

The metabolic energy cost for human walking varies considerably with terrain. For example, loose sand can double the cost compared to a smooth, hard surface [1,2]. Overall energy expenditure is also determined by other variables such as carried load, movement speed, and grade or ground slope [3–5], each with readily identifiable effects. But the effect of terrain could depend on more complex factors such as unevenness of the surface, its compliance and energy absorbing properties, and looseness and instability of the substrate. That complexity is typically avoided in predictions of metabolic cost, in favor of a single multiplicative factor, the *terrain coefficient*, for the relative gross metabolic cost compared to treadmill walking. Typical values are 1.0 for blacktop surface, 1.2 for light brush, 1.5 for heavy brush, and 2.1 for loose sand [2]. But aside from this overall effect, there is presently scant understanding of how terrain affects a person’s actual movements and actions, which are the ultimate determinants of energy expenditure. If the gait adaptations for different terrains could be quantified, they might offer insight regarding the control of locomotion and improved predictions for its energetic cost.

It is challenging to determine the biomechanical adaptations for different terrains. Traditional laboratory measures include kinematics and ground reaction forces [6], which can yield mechanistic measures such as fluctuations in kinetic energy when walking on sand [1], or the work performed by the leg joints on an artificial, uneven treadmill surface [7], with attendant energetic cost. But such laboratory measures are difficult to obtain outdoors. This limitation favors simpler equipment such as body-worn accelerometers, whose signals can be correlated with energy expenditure [e.g., 8–12], albeit with limited ability to distinguish terrain type [13]. Yet another possibility is to use shoe-mounted inertial measurement units (combining accelerometers and gyroscopes) to reconstruct the foot’s path in space and placement on ground [14,15]. These data can reveal trends in walking speed, stride length, and stride variability [16], which may in turn reveal the effects of real-world terrain.

Ground terrain could have various effects on the foot’s motion during walking. Most obvious is the elevation change over a step, which is energetically costly for a net elevation increase [17], and might also increase cost for terrain that undulates from step to step with no overall slope. Terrain might also affect parameters such as average stride length and width, which also determine energy expenditure [e.g., 18,19]. Uneven terrain may require the foot to be lifted higher mid-swing [20], with an attendant cost [21]. Finally, balance might be more challenging on some terrains, requiring stabilizing adjustments [22] including foot placement [7,23]. Thus, motion of the foot may entail energy expenditure.

The purpose of this study was to determine how foot paths change with terrain, and how they relate to the energetic cost of walking. Here, *foot path* refers to the foot’s translation in three dimensions during a single swing phase, starting from the previous stance phase and including the ending stance phase, when the foot is stationary. We tested whether this path exhibits changes in standard gait measures, such as average stride length and height and their respective variabilities, as a function of terrain. We also tested these measures for correlation with energy expenditure, to examine the possible link between foot path, energy cost, and terrain.

## Methods

We measured healthy adults walking on five types of common outdoor surfaces: Sidewalk, Dirt, Gravel, Grass, and Woodchips (see Figure 1). The experiment was performed outdoors in Nichols Arboretum (Ann Arbor, MI), a University-operated park with well-groomed walking trails, selected to pose little challenge to any healthy individual. For all conditions, subjects followed trails intended for walking, except for Grass which was in a meadow without a specific trail. All of the surfaces were selected to have very little elevation change, in terms of visible undulations, total change (maximum net grade of 0.96% on Gravel), and cross-slope. We measured metabolic energy expenditure, foot paths, and attendant stride parameters during walking. Stride information was collected using inertial measurement units (IMU) (Opal sensors, APDM Inc., Portland, OR) attached atop each foot. A global positioning system device (GPS; Garmin Ltd., Olathe, KS) was also used to characterize the route’s speed, distance, and elevation.

**Figure 1.**
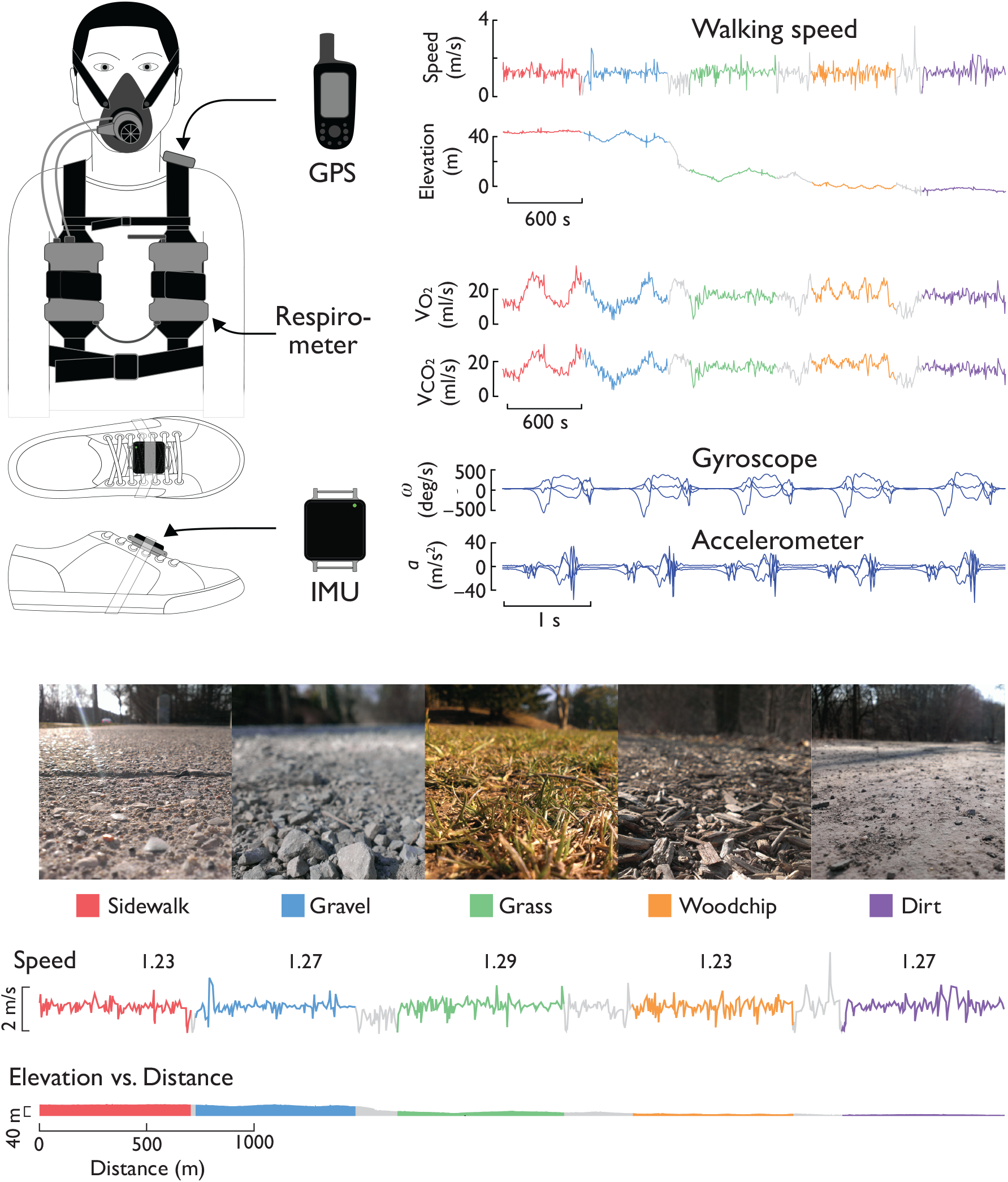
Measurement of foot paths and energy expenditure on outdoor terrain. Subjects walked on different terrains while wearing a portable respirometry system, a global positioning system (GPS) device, and one inertial measurement unit (IMU) per foot. Sample data from one subject show traces for walking speed and elevation from GPS, rates of oxygen consumption and carbon dioxide production, and angular velocity and translational acceleration vs. time. Terrains included Sidewalk, Gravel, Grass, Woodchip, and Dirt, along with transitions between them (gray lines, not analyzed). Walking speed was loosely regulated via GPS (average speeds listed); terrain segments were selected to avoid large net changes in elevation during trials.

## Experiment

Ten adult subjects (N=10, 5 male and 5 female, age 18 - 48) participated in the study. Subjects had an average body mass of 64.86±10.10 kg (mean ± s.d.) and an average leg length of 0.90±.07 m (mean ± s.d.). Subjects provided written informed consent before the experiment. The study was approved by the University of Michigan Health Sciences Institutional Review Board (HUM00020554).

Subjects walked on each surface, presented in random order, for 8 minutes. Approximate speed of 1.25 m/s was controlled by following the experimenter, who walked according to GPS speed and attempted to make only gentle speed corrections, to avoid costs for artificial speed fluctuations [24]. Some surfaces were limited in length, and so subjects reversed their direction and continued walking. Turns occurred at most 10 times per 8-minute trial.

Respirometry data were collected for the entirety of each trial (Oxycon Mobile, CareFusion Corp., San Diego, CA). To allow time to reach steady-state, only the last 3 minutes of data from each surface were used for metabolic energy expenditure. The rates of oxygen consumption and carbon dioxide production (mL/min) were converted to metabolic rate (W) using standard formulae [25,26]. Net metabolic rate *Ė*_met_ was calculated by subtracting metabolic rate of a separate quiet standing trial (97.29 ± 27.06 W) from gross. We also calculated a dimensionless net metabolic cost of transport, defined as the net energy expended to move a unit body weight a unit distance.

For each trial, a total of 90 strides per foot were analyzed from forward walking sections at the beginning of the trial (Figure 2). Estimated foot paths were derived from IMU data according to an algorithm described previously [15]. Briefly, the method uses gyroscope and accelerometer data to estimate spatial orientation, and then integrates translational accelerations twice to yield displacements, with inertial drift reduced by correcting the velocities during stance to zero. Here, foot path actually refers to the path of the IMU, located on the instep of the shoe. From these paths, we computed gait parameters such as stride length, width, and height, all defined as displacements over one stride. To reduce the amount of data, only the left foot data were used for the measures reported here. We report average and root-mean-square (RMS, equivalent to standard deviation) variability of stride parameters, except for average stride width, which was unknown because each IMU recorded independent data for one foot, with no reference to the other foot. We also estimated two additional parameters defined by the foot’s stationary positions at beginning and end of stride, and the straight line connecting those positions. Projected onto the sagittal plane, the *virtual clearance* was defined as the closest distance the foot reaches to this line (measured perpendicularly) during the middle of swing phase (illustrated in Figure 2), extending a measure previously defined for flat ground [27] to include different footfall heights. Projected onto the transverse plane, *lateral swing displacement* was defined as the maximum distance the foot departs from this line, also mid-way through the swing phase.

**Figure 2.**
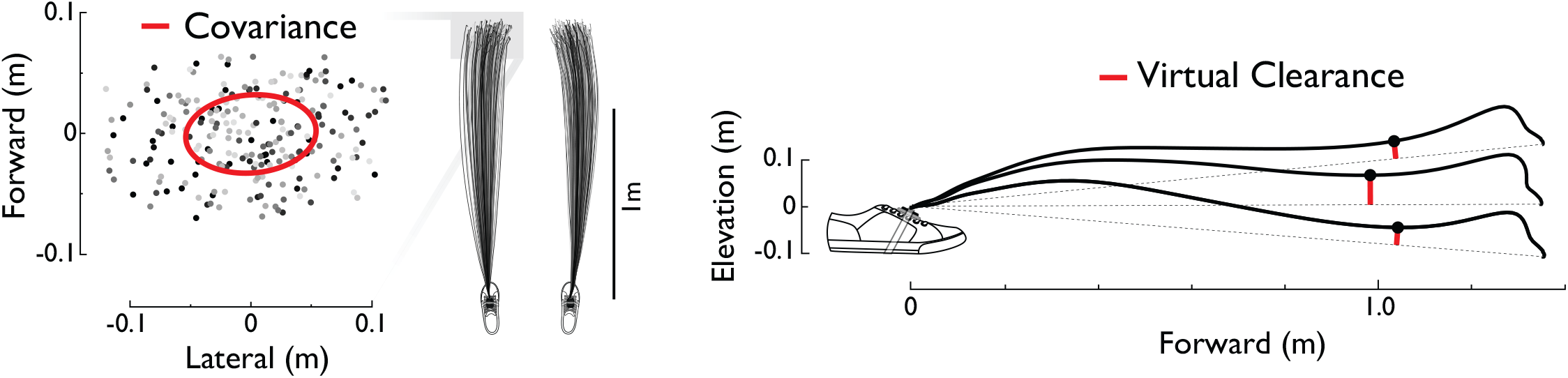
Sample foot path trajectories and associated measurements, as viewed from above and from the side. Forward vs. lateral foot displacements from each trial were used to compute stride covariances. Vertical path of foot was used to determine virtual clearance, relative to straight line between start and end of stride.

Stride parameters and energy measures were normalized to account for differences in subject body size and height. We used body mass *M*, standing leg length *L* (defined as floor to greater trochanter), and gravitational acceleration *g* as base units. Thus, stride distances were normalized by *L*, and net metabolic power [28] by *Mg*^1.5^*L*^0.5^ (average 0.90 m, 1893 W across subjects). Quantities were then reported in dimensional form by multiplying by the mean normalization factor across subjects.

We tested whether terrain conditions affected energy expenditure and gait parameters. We calculated the mean and standard deviation of the measures across subjects for each terrain surface. Differences between the conditions were quantified by repeated-measures ANOVA tests. We also tested the correlation between energy expenditure and the gait parameters using linear regression for each variable individually. The latter included a separate offset constant for each individual, included in the fit, with overall goodness of fit therefore evaluated with an adjusted *R*^2^. The significance level *α* was set at 0.05.

To explore reduction of dimensionality within the data, we also performed principal components analysis (PCA) and linear discriminant analysis (LDA). The PCA was intended to reduce the 11-dimensional stride measures into a smaller number of combinations, and reveal which combinations contribute most to the observed variations, without regard to terrain type. The LDA (using only linear terms for each predictor) was performed to use the same data to classify the terrains, with knowledge of each trial’s terrain included. Finally, an additional set of regressions was performed between metabolic rate and stride measures, using principal components regression (PCR) and partial least squares regression (PLSR), to determine how a small set of data combinations can predict metabolic rate, again with adjusted *R*^2^ to evaluate goodness of fit.

## Results

We found the foot paths to be highly dependent on terrain. This was observable qualitatively in the foot paths, which showed changes in variability compared to the Sidewalk condition as viewed from the side and above (see Figure 3 for representative paths). Such terrain-related differences were also confirmed quantitatively for most of the stride parameters considered (Figure 4), particularly the measures of virtual clearance (mean changing by up to 58% and variability by up to 63%), and to lesser degree, lateral swing displacement (mean and variability, summarized in Table 1).

**Table 1.** Stride measures and energy expenditure for five terrains. Results are shown as mean ± s.d. across subjects (*N* = 10). Significance (S) of each measure indicated by asterisk ‘*’ (repeated measures ANOVA, *P* < 0.05).

| Measure |  | Sidewalk | Dirt | Gravel | Grass | Woodchips | S | P |
| --- | --- | --- | --- | --- | --- | --- | --- | --- |
| Virtual Clearance (m) | Mean | 0.031±0.008 | 0.033±0.009 | 0.041±0.009 | 0.049±0.008 | 0.050±0.009 | * | 4.66e-08 |
| | RMS | 0.006±0.001 | 0.006±0.002 | 0.007±0.001 | 0.010±0.002 | 0.011±0.003 | * | 3.70e-04 |
| Lateral Swing (m) | Mean | 0.039±0.015 | 0.040±0.013 | 0.043±0.016 | 0.039±0.012 | 0.043±0.016 | * | 1.28e-12 |
| | RMS | 0.016±0.003 | 0.017±0.003 | 0.019±0.003 | 0.019±0.004 | 0.020±0.004 | * | 6.69e-05 |
| Stride Height (m) | Mean | -0.004±0.021 | -0.013±0.019 | 0.056±0.027 | 0.038±0.028 | 0.019±0.062 | | 1.74e-01 |
| | RMS | 0.014±0.002 | 0.016±0.004 | 0.028±0.003 | 0.031±0.003 | 0.044±0.008 | | 2.84e-01 |
| Stride Length (m) | Mean | 1.411±0.069 | 1.470±0.064 | 1.404±0.096 | 1.440±0.064 | 1.451±0.054 | * | 1.84e-09 |
| | RMS | 0.033±0.005 | 0.027±0.005 | 0.034±0.005 | 0.038±0.003 | 0.043±0.005 | | 6.35e-02 |
| Stride Width (m) | RMS | 0.051±0.009 | 0.056±0.006 | 0.066±0.010 | 0.068±0.008 | 0.099±0.013 | | 4.23e-01 |
| Speed (m/s) | Mean | 1.281±0.085 | 1.334±0.089 | 1.263±0.118 | 1.279±0.085 | 1.287±0.085 | * | 1.04e-13 |
| | RMS | 0.048±0.009 | 0.038±0.006 | 0.047±0.007 | 0.050±0.004 | 0.059±0.009 | * | 1.53e-04 |
| Net Metabolic Rate (W) | Mean | 189.1±29.00 | 204.4±35.96 | 218.9±35.62 | 223.2±28.27 | 240.8±28.91 | * | 7.11e-11 |
| Net Cost of Transport | Mean | 0.232±0.036 | 0.241±0.038 | 0.272±0.033 | 0.275±0.035 | 0.294±0.031 | * | 3.28e-09 |

**Figure 3.**
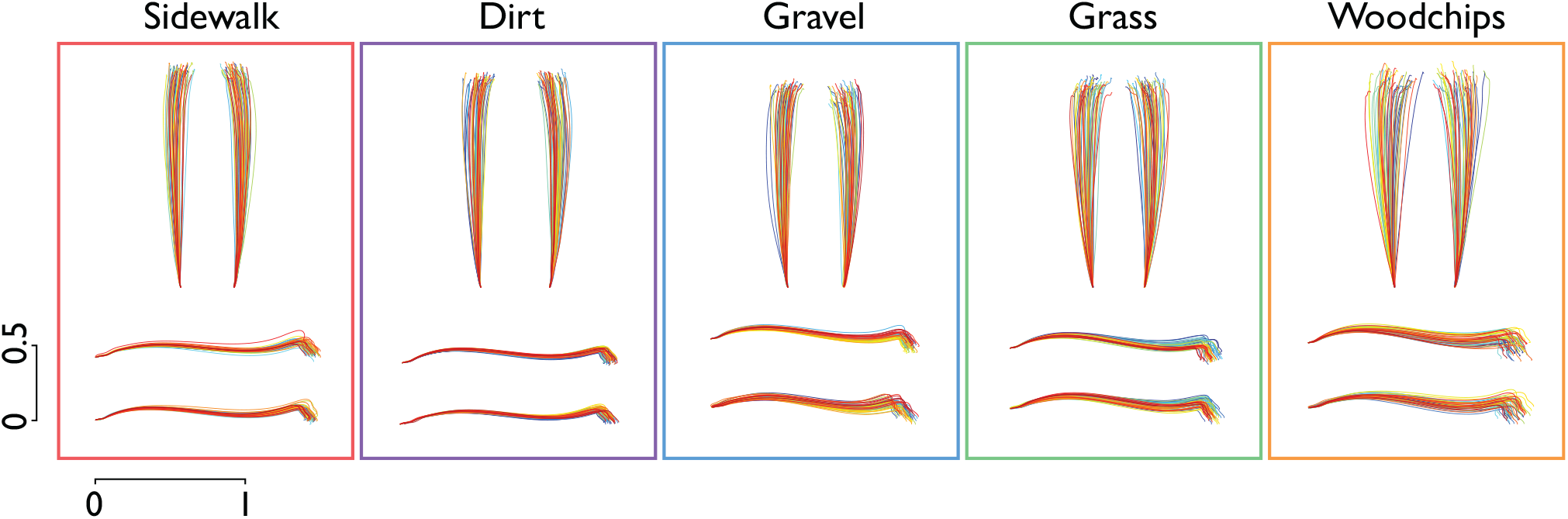
Representative foot path trajectories for each terrain (from one representative subject), as viewed from above and from side. All strides were arranged to have common origin, to emphasize variation among strides. Color of trajectories varies gradually between beginning (blue) and end (red) of trial, to indicate time course of strides.

**Figure 4.**
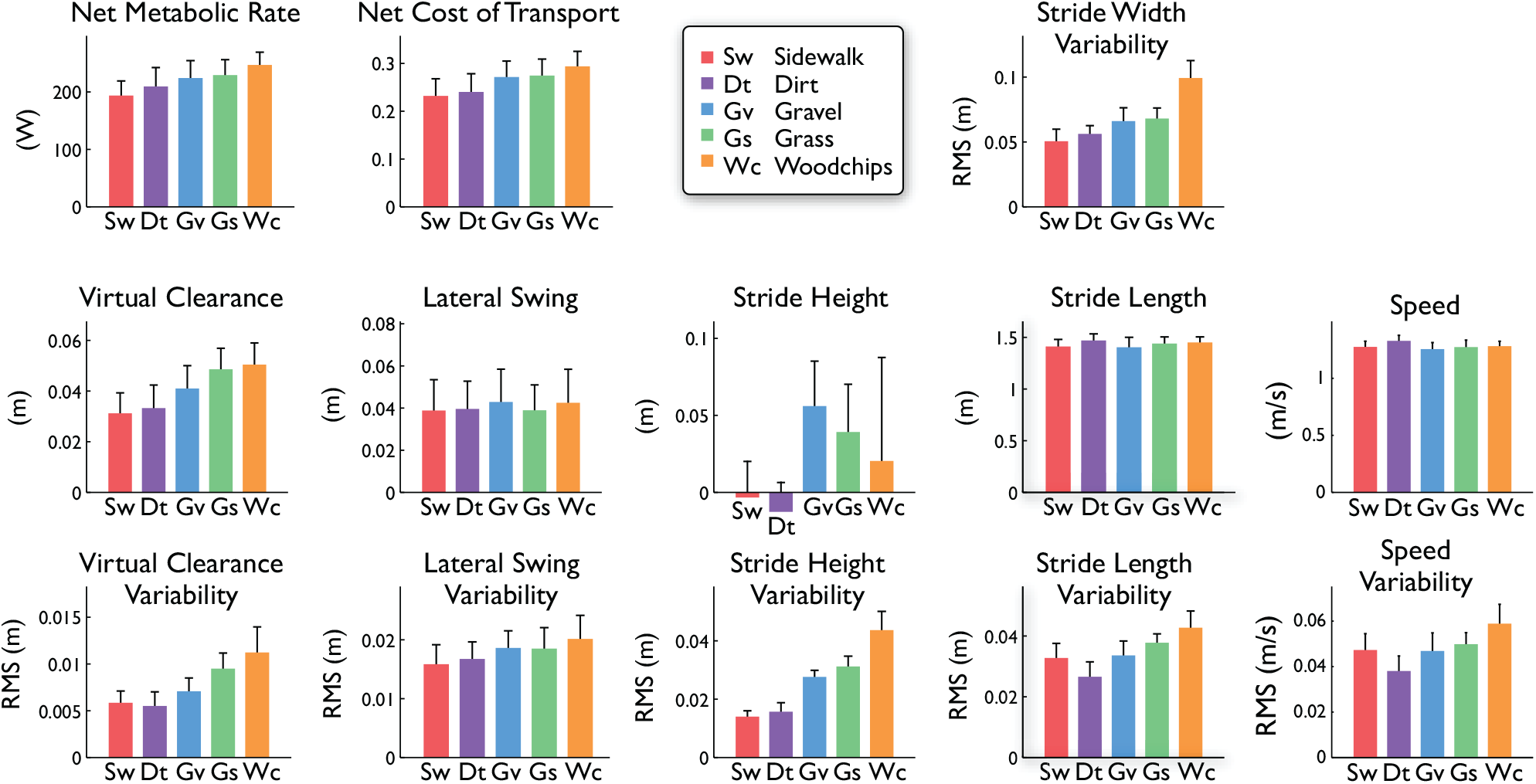
Summary measures of energetic cost and stride measures on five different terrains. Energy expenditure in terms of net metabolic rate and net metabolic cost of transport (energy per unit distance and weight). Stride measures are shown as mean and root-mean-square (RMS) variability: virtual clearance, lateral swing distance, stride height, stride length, stride width (variability only), and walking speed. Bars denote across-subject means; error bars denote standard deviation across subjects (*N* = 10).

Participants also expended varying amounts of energy as a function of terrain (Figure 4, top). Net metabolic rate *Ė*_met_ varied with terrain type for groupwise (repeated measures ANOVA, *P* = 7.1e-11) and for most pair-wise comparisons (post hoc paired t-tests, *P* < 0.05), with the greatest difference (27%) found between Woodchips and Sidewalk. The only non-significant comparisons were Dirt vs. Sidewalk, Gravel vs. Dirt, and Grass vs. Gravel (*P* ≥ 0.05). Summary results below are presented in order of increasing mean metabolic rate: Sidewalk, Dirt, Gravel, Grass, Woodchips.

Stride parameters also correlated with metabolic rate irrespective of terrain classification. From linear regression, nearly every stride measure was found to be significantly correlated to metabolic rate *Ė*_met_ (Table 2); the only non-significant measures (*P* ≥ 0.05) were mean walking speed and lateral swing (mean and variability). For goodness of fit, the top four correlates were mean virtual clearance, and RMS variabilities of virtual clearance, stride height, and stride width. These measures were all strongly significant regressors (at most *P* =3.1E-06), although the actual predictive abiilty was modest, with adjusted *R*^2^ ranging 0.29 – 0.38). Part of the variation within the data may be attributed to inter-subject differences. This was revealed by improved fits (Table 2, “Ind *R*^2^”)when subject-specific offsets were removed from metabolic data, yielding for example an increase of 0.15 (i.e. a partial *R*^2^)for mean virtual clearance.

**Table 2.** Linear relationship between net metabolic rate (outcome variable) and individual stride measures. Linear regression was performed on each measure, yielding a slope (with 95% confidence intervals, c.i.) and constant offset, as well as adjusted *R*^2^ and individualized adjusted *R*^2^ (with separate offset for each subject, “Ind”). The difference between individualized and traditional *R*^2^ indicates how much of the variability was due to subject offsets, as opposed to terrain type. Significance (*P* < 0.05) of regression indicated by dagger ‘†’, and significant difference in regressor across terrains by asterisk ‘*’ (identical to Table 1). Regression slopes are reported in units of W/m for all regressors except speed (W · s · m^−1^), and offsets in units of W.

| Regressor | | Slope | ± | c.i. | Offset | $R^2$ | Ind $R^2$ | S | P |
| --- | --- | --- | --- | --- | --- | --- | --- | --- | --- |
| Virtual clearance | Mean | 1714. | ± | 399.9 | 155.6 | 0.34 | 0.49 | †* | 2.63e-11 |
| | RMS | 5948. | ± | 1657. | 182.5 | 0.29 | 0.46 | †* | 3.4e-09 |
| Lateral swing | Mean | 89.05 | ± | 494. | 224.6 | -0.01 | 0.00 | * | 0.719 |
| | RMS | 2329. | ± | 1729. | 187.4 | 0.06 | 0.07 | †* | 0.00937 |
| Stride height | Mean | 167.3 | ± | 139.1 | 225.1 | -0.02 | 0.08 | † | 0.0195 |
| | RMS | 1410. | ± | 353.6 | 193.1 | 0.30 | 0.56 | † | 2.05e-10 |
| Stride length | Mean | 88.77 | ± | 86.88 | 102.3 | 0.38 | 0.03 | †* | 0.0454 |
| | RMS | 1414. | ± | 820.2 | 174.3 | 0.02 | 0.18 | † | 0.00113 |
| Stride width | RMS | 689.3 | ± | 262.6 | 185.1 | 0.14 | 0.35 | † | 3.11e-06 |
| Speed | Mean | 1.96 | ± | 68.6 | 226.3 | 0.19 | 0.00 | * | 0.954 |
| | RMS | 1067. | ± | 593.3 | 170.1 | 0.02 | 0.16 | †* | 0.000713 |

Principal components analysis revealed that the first two PCs could explain a substantial fraction of the observed stride measures (Figure 5). The first PC accounted for 65.8% of all terrain-specific variability in the stride measures, and was dominated by increased stride length, increased walking speed, and negative stride height (apparent downhill slope). The second PC accounted for an additional 21.7% (and thus both PCs 87.5%), and was dominated by increased stride length, increased stride height (apparent uphill slope), and increased stride width variability. These two PCs (together accounting for 87.5% of all data variability) were subsequently used as regressors of metabolic rate.

**Figure 5.**
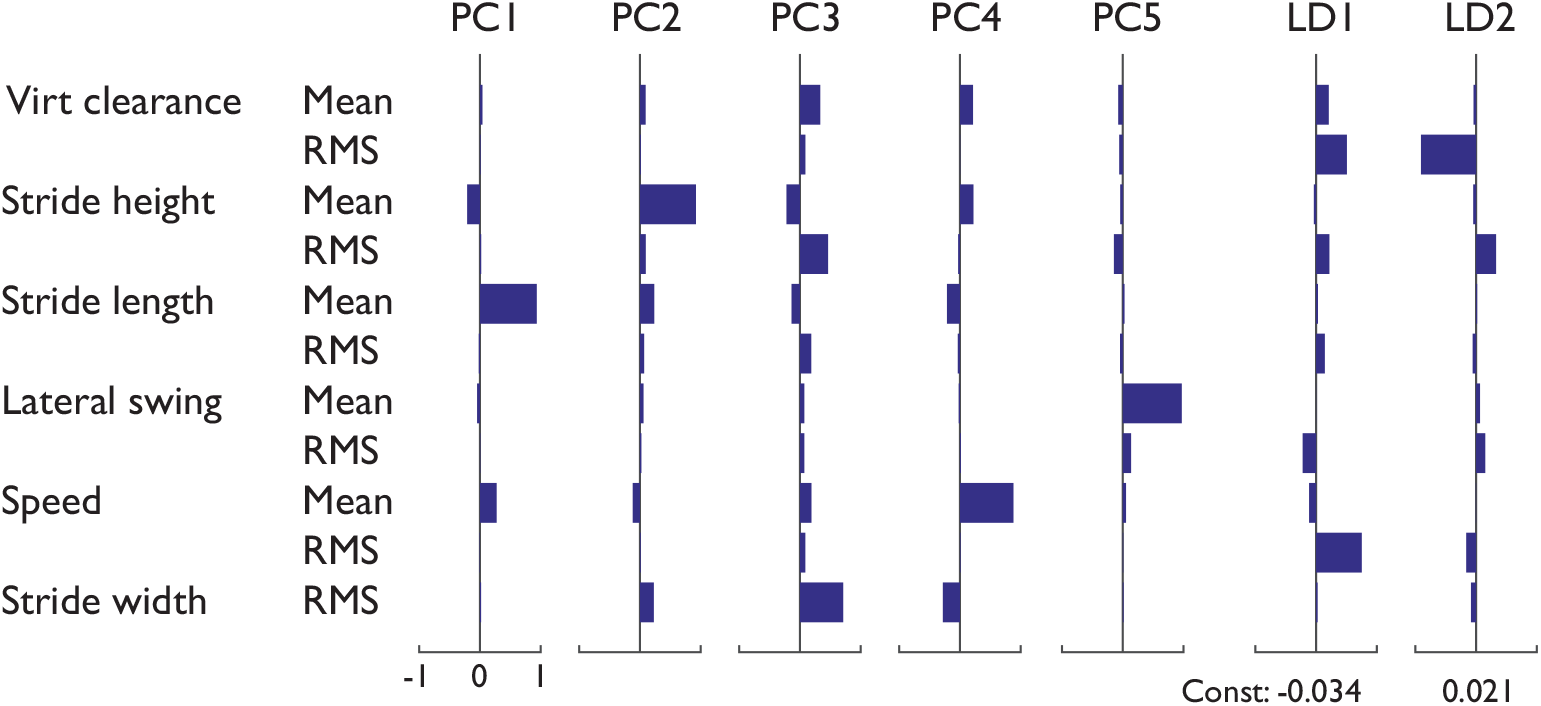
Principal components and linear discriminants of stride measures, shown as a series of columns of horizontal bars, each row representing a stride measure. First five principal components (PCs) are shown, as well as two linear discriminants, for (LD1) Gravel vs. Grass, and (LD2) Sidewalk vs. Dirt (with constant offsets listed). Stride measures from all subjects and all terrains contributed to this analysis.

Linear discriminants were able to classify the data reasonably well (Figure 5), with 10% resubstitution error rate (5 errors out of 50 observations from 5 terrains and 10 subjects). This was true despite substantial overlap between terrains and subjects in individual measures such as stride length vs. speed (Fig. 6, top). To illustrate the classification, we projected the stride measure data onto two sample discriminants: Gravel vs. Grass, and Sidewalk vs. Dirt, two pairs poorly distinguished by the individual stride measures. The discriminated data (Fig. 6, bottom) show reasonably good discrimination between those same pairs.

**Fig. 6.**
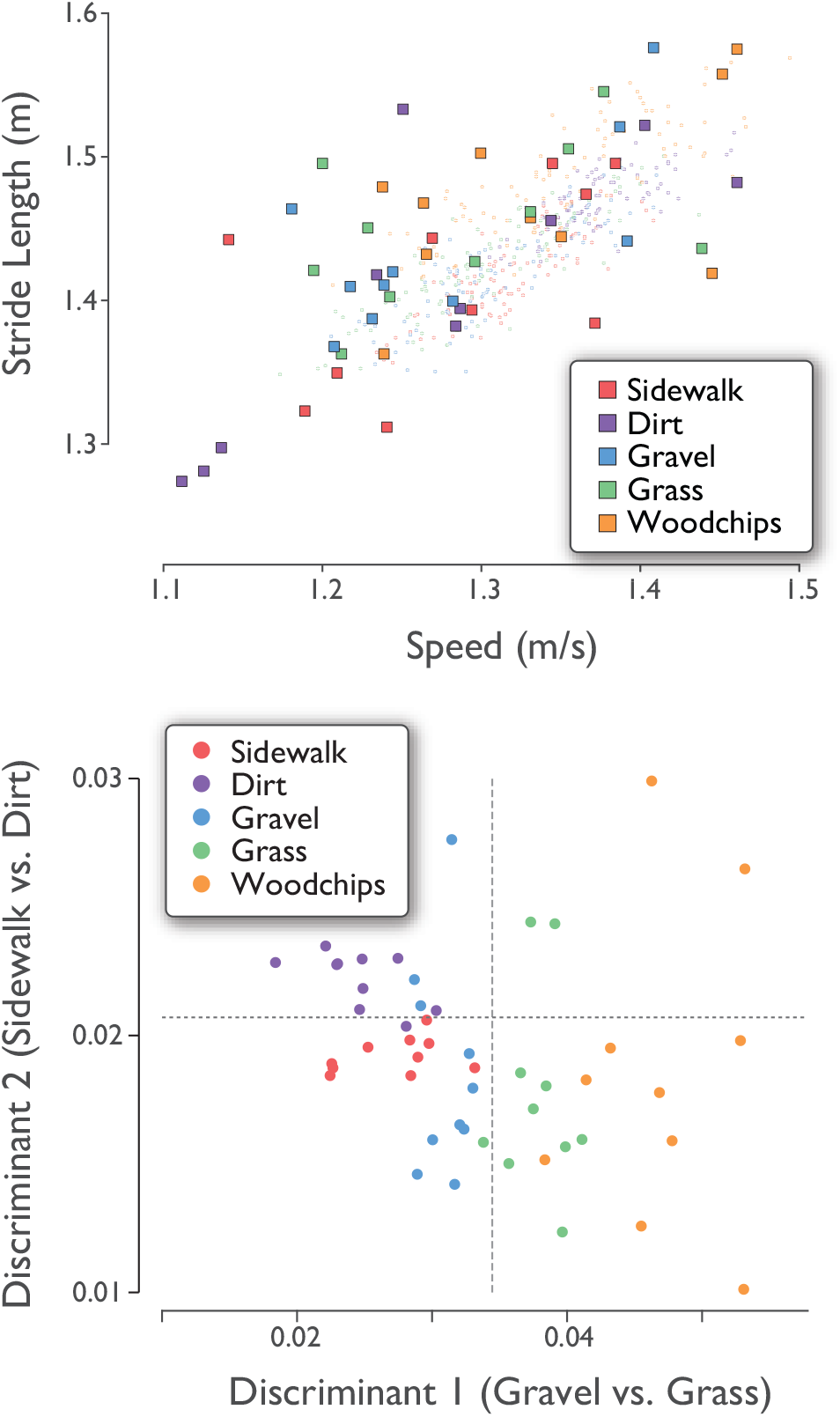
Stride measures for all subjects (*N* = 10) and all terrains, plotted in two ways: (top) Stride length vs. Speed, and (bottom) Linear discriminants against each other (i.e. a projection of multi-dimensional data onto two discriminants). Each data point represents one subject’s average measures for one terrain. Stride lengths and speeds (filled symbols) were highly correlated with each other, and overlapped for different terrains. As an example of within-trial variations, top graph also shows all strides from all terrains for a single representative subject (smaller, lightly shaded symbols). Linear discriminants improve separation between two pairs of terrains (separators denoted by dashed lines).

Although we attempted to approximately control the average walking speed, there was some variation within each trial. Walking speed normally fluctuates slightly [29], with correlated fluctuations in stride length [16] consistent with the preferred stride length relationship [30]. Some individuals exhibited terrain-dependence in their relationship (Figure 6, top), but with no consistent statistical trend across subjects. Thus, the preferred stride length vs. speed relationship remained fairly intact across different terrains. There were also small but significant differences in mean speed and stride length across terrains (Table 1).

Metabolic rate was explained reasonably well with all three methods considered (Fig. 7). The best explanation resulted from partial least squares regression (PLSR), which uses all stride measures and metabolic outcome data together to define a set of multivariate regressors (defined in Table 2). This technique yielded adjusted *R*^2^ = 0.52 to predict metabolic rate using only two such regressor components. In contrast, principal components regression (PCR) first derives principal components to explain variations within the stride measure data (without considering outcome data), and then uses those components for regression. Using only the first two PCs (described above), PCR yielded *R*^2^ = 0.46 (see Table 2). Both of these exceed the fit for the strongest single univariate regression (virtual clearance, with *R*^2^ = 0.34). As few as two multivariate regressors can therefore explain a greater proportion of the variations in the outcome data, compared to any single measure.

**Figure 7.**
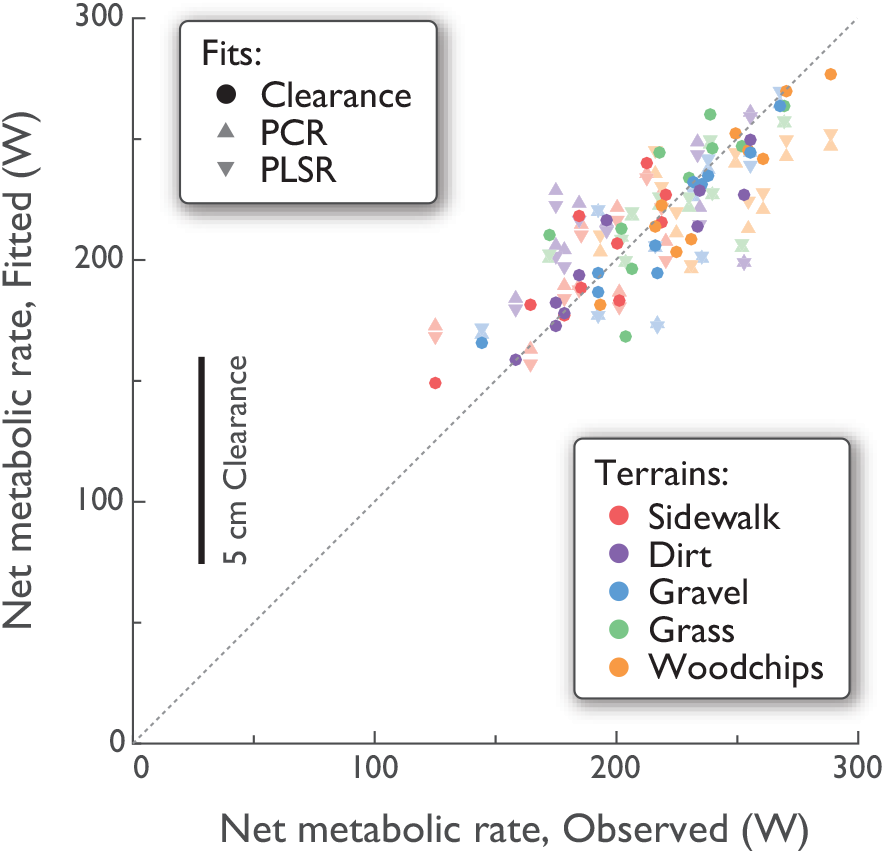
Net metabolic rate for all subjects and all terrains, fitted vs. observed. Observed refers to empirical measurements (five terrains, *N* = 10 each). Fitted refers to three ways to predict metabolic rate: Principal components regression from first two PCs (PCR; adjusted *R*^2^ = 0.46); Partial least squares regression (PLSR; adjusted *R*^2^ = 0.52; and from virtual clearance in a single-variable linear regression (Clearance; overall adjusted *R*^2^ = 0.34; shown fitted with subject-specific offsets, *R*^2^ = 0.49). Fit types are denoted by symbol shape, and terrains by color.

## Discussion

This study tested for relationships among the foot’s path and placement, the type of ground terrain, and the energy expended for walking. We found that multiple stride parameters are indeed terrain-dependent and correlated with energy cost. Notably, more challenging terrain caused increases in virtual ground clearance and in the variability of most measures, for example of lateral swing motion. These measures were in turn correlated with increased energy cost. Any single measure could only predict metabolic rate imperfectly, but there was also considerable interdependency among measures, as revealed by dimensionality reduction techniques. We found that both principal components analysis and partial least squares regression could yield reasonable predictions of metabolic cost based on as little as two multivariate components. We next provide our interpretation of the relationship between stride measures and metabolic cost on different terrains, and their possible utility.

Participants made only subtle changes to their average gait pattern as a function of terrain. Most notable was virtual clearance of the swing foot, which increased on more challenging terrain (Table 1), and was highly correlated with energy expenditure (Table 2). The latter is consistent with controlled experiments showing a high cost for increased clearance [21]. Of course, the details of actual surface variations were unknown, and so virtual clearance is merely an indicator of possible adaptations to true ground clearance. There were also small changes in stride length and speed with terrain, which may be attributable in part to imperfectly controlled walking speed rather than the terrain itself.

While the average gait pattern changed little, variability in most of the gait measures examined showed high dependence on terrain. The most notable sensitivities were for variability in stride height, stride width, virtual clearance, and lateral swing motion. Variability could result directly from the unevenness of ground, or from controlled adjustments made to stabilize balance, which is thought to be passively unstable in the lateral direction [22,23]. Active stabilization is achieved in part through lateral foot placement [23,31–34]. Uneven ground appears to disrupt gait to substantial degree, and would be expected to require substantial active stabilization. Aggregating these various contributions, the overall effect is that uneven ground leads to uneven foot motion and uneven steps.

Stride measures also appear to be predictive of energy expenditure. Nearly every stride measure exhibited significant correlation with energy expenditure, most strongly the RMS variabilities of stride height, virtual clearance, and stride width (Table 2). Walking speed is generally a strong predictor of energy cost [5,35]. Our interest here was in factors other than speed, which we therefore attempted to control at fixed value across terrains (e.g. 0.5% speed difference between Woodchips and Sidewalk). Thus, the weak and non-significant correlation between speed and energy cost (Table 2) was merely a consequence of experimental control rather than a finding. Walking speed also generally determines stride length [16,29,36], which was not explicitly controlled and differed slightly with terrain. By itself, stride length was a barely significant correlate of energy cost (Table 2), which could be due in part to an actual effect, and in part to imperfect experimental control of speed. Indeed, co-variation of speed and stride length dominated the first principal component of stride measures (Fig. 5), and predicted energy expenditure from the principal components regression (PCR, Fig. 7). In addition, all stride variability measures were individually correlated with energy cost (Table 2), although they contributed relatively little to the first two principal components. Variability in stride length and timing [37] and fluctuations in speed [24] have been reported to affect metabolic cost, perhaps due to the effort of varying gait. These results illustrate the importance of interdependencies among stride parameters, and the complex relationship of cost to gait parameters.

Another well-known predictor of energy expenditure is elevation change. Even though elevation changes were modest on the terrains studied here, a non-zero stride height would generally be expected to indicate how much the body is lifted or lowered against gravity, and therefore drive energy expenditure. Other cost-determining variables more specific to terrain included virtual clearance and its variability, and variability of stride height and width. If a single predictor is desired that is both sensitive to terrain and predictive of energy expenditure, the strongest candiate is virtual clearance (Fig. 7), followed by lateral swing variability, which may be an indicator of the balancing challenges posed by uneven ground.

Alternatively, the PCR and PSLR results show that IMU-derived foot paths can also yield multivariate components, or linear combinations of measures, that can be more reliably predictive than any single variable. Of course, IMU-based measures are unlikely to replicate the accuracy of a (portable) respirometry system, but IMUs are less obtrusive and easier to wear, especially in real-world conditions, and may still yield data informative of metabolic cost.

Stride measures may also serve as a supplement to terrain classification. A terrain such as “grass” can vary substantially in height, thickness, density, and underlying substrate, which itself may vary in softness, granularity, friction, and moisture content. Even if terrain were accurately imaged and quantified for geometric scale and irregularity [38], there may be a plethora of variables relevant to gait. In contrast, a few stride measures, such as stride and swing foot variability (Figs. 5 and 6) can directly measure a terrain’s effect on gait, and even discriminate among terrains. Gait measures are unlikely to discriminate better than visual observation, but they do offer continuous quantification of a terrain’s effects. Just as the classification of “highway” might be supplemented by information about traffic and road conditions, a prospective hiker or trekker might gain from knowledge of a “grass” trail’s typical effects on stride variability, time to destination, or metabolic cost (Fig. 7). There may well be benefit to quantifying terrain by entire new continuous measures or discrete categorizations, independent of semantic classifications.

This work is subject to a number of limitations. We based our analysis on a relatively small number of summary measures, but a more intensive approach might be to instead use the actual foot path trajectories directly, including both translation and orientation data. The much larger volume of source data, with appropriate data reduction, might yield stronger classifiers and correlators. Another limitation of the present foot path reconstruction technique is that measurement errors are unavoidably greater than those typical for laboratory motion capture. Our foot path estimation relies on the foot being nearly stationary at some point during stance, which may not occur for every stride on softer terrains such as Woodchips.

This adds significant uncertainty to estimates of stride height and its variability in these conditions. Indeed, all of the variability reported here is in part due to terrain, inertial drift, and other measurement noise, in addition to true motion variability. In particular, there can be vast variations between terriains of a single type such as Sidewalk. Each location in the world, whatever its classification, may have unique effects on gait, that may nonetheless be quantifiable.

There are also limitations to the degree that kinematic measures can explain energy expenditure. Energy cost depends considerably on mechanical work performed by the body [39], even on uneven terrain [7], but foot paths cannot capture the force or power produced by the leg. In addition, inertial data cannot readily discern step width, which also appears to change on uneven terrain [7] and could contribute to energy cost [18]. Thus, IMU-derived foot paths are neither absolute nor comprehensive measures. More complete kinematic data are obtainable with IMU suits (e.g., Perception Neuron suit, Noitom Ltd, Miami FL USA), which might improve upon our results. We find that foot-mounted IMUs appropriately meet the trade-off between data quantity and convenience and practicality for real-world usage.

An improved study would include more variables than examined here. This could include more challenging terrain with significant speed and elevation variations, or with carried loads, to evaluate the interactions that determine energy expenditure [5,40,41]. Measures of gait and energy expenditure could conceivably be combined with geographical information systems (GIS) technology and embedded into map databases [42]. Although foot motion hardly encompasses all of the gait adaptations for terrain, it is highly sensitive to the type of terrain, and has a discrete ability to categorize or discriminate terrains objectively. It also exhibits a continuous correlation with energy expenditure, which could potentially have predictive applications.

## Acknowledgements

This work was supported in part by Department of Defense (W81XWH-09-2-0142), National Institutes of Health (AG030815), and Office of Naval Research (ETOWL).

